# Phenogenon: Gene to Phenotype Associations for Rare Genetic Diseases

**DOI:** 10.1101/367292

**Authors:** Cian Murphy, Ismail Moghul, Nikolas Pontikos, Phenopolis consortium, UK Inherited Retinal Dystrophy consortium, UCLex consortium, Jing Yu

## Abstract

As genome sequencing is increasingly applied to molecular diagnosis of rare Mendelian disorders, large number of patients with diverse phenotypes have their genomic and phenotypic data pooled together to uncover new genotype - phenotype relations. We introduce Phenogenon, a method that combines: the power of Human Phenotype Ontology for describing patient phenotypes, gnomAD for estimating rare variant population frequency, and CADD for variant pathogenicity prediction. By using a divide and conquer approach, we demonstrate here that Phenogenon is able to uncover true gene to phenotype relations, such as *“ABCA4* – Macular dystrophy” and *“SCN1A* – Seizures”. Additionally, it accurately infers mode of inheritance, such as *a* recessive mode of inheritance in the case of the “*ABCA4* – Macular dystrophy” relationship and *a* dominant mode of inheritance with the “*SCN1A* – Seizures” relationship. We also found that CADD has more power to detect early-onset rare genetic diseases than late-onset diseases. In this study, we ran Phenogenon against a diverse cohort of 3288 patients. Among the top 13 gene-phenotype relations, seven were previously known. We also highlight four potentially novel gene – phenotype relations such as “*SIPA1L3* – Abnormal electroretinogram”.

## Introduction

As DNA sequencing cost decreases, whole exome sequencing (WES) that covers almost all protein coding regions of the genome has gained popularity in the molecular testing of individuals with rare Mendelian disorders. The resultant data deluge of variants of unknown pathogenicity and clinical significance is an onerous task to interpret. To help prioritise, a common practice is to search similar disease cases for shared genetic variations that are already solved. Conventionally, this is done by checking published literature and databases such as dbSNP^1^ and ClinVar^2^ for genetic variants and Online Mendelian Inheritance in Man (OMIM), for genes. For well-characterised diseases such as retinal dystrophy and epilepsy, one can also use targeted databases such as RetNet (https://sph.uth.edu/Retnet/) and EpilepsyGene (http://www.wzgenomics.cn/EpilepsyGene/), respectively. Despite this, it is common that no candidate genes or variants are found in published solved cases. Therefore, one solution is to group unsolved cases with similar phenotypes in order to increase the chance of finding shared genetic variations.

The UK Inherited Retinal Dystrophy Consortium (UKIRDC) adopted this initiative by collecting unsolved retinal dystrophy patients from University College London, University of Oxford, University of Leeds and University of Manchester. At the time of writing, 365 exomes have been collated. The exome and phenotype data are deposited as part of the UCL Exome Consortium (UCLex), which itself hosts 3288 of unrelated exomes and consists of patients with a range of disorders such as dementia, Crohn’s disease, seizures and bone-marrow failure (Supplementary Fig. 1)^3^.

Nonetheless, having to sift through a large number of samples for candidate genetic variations that are enriched only in patients with similar phenotypes remains a daunting task. Here we propose Phenogenon, which evaluates gene-phenotype associations from large and mixed cohorts of patients. Phenogenon categorises genetic variants according to their population frequencies and predicted pathogenicity, and reports the association between patient carriers of each group and a given phenotype. An overall **H**PO **G**oodness of **F**it (HGF) score can be calculated for each gene and phenotype by combining the statistics of the grouped analyses. In this manner, the HGF score allows for easy prioritisation of genes for a given phenotype. For instance, when analysing the UCLex dataset with Phenogenon, we showed that the majority of the ‘Seizures’ cases were associated with a single mutation in *SCN1A*, and in the case of ‘Retinal dystrophy’, two mutations in the *USH2A gene*. Conversely, we were also able to infer the strongest associated phenotype of a given gene, such as ‘Macular dystrophy’ for *ABCA4*. Additionally, Phenogenon is able to provide an indication of the mode of inheritance of a gene-phenotype relation.with reasonable accuracy.

## Materials and methods

### Data collection

The patient phenotypes which contributed to this project came from the UCLex which currently includes 5122 Mendelian and common disease patients. We used Human Phenotype Ontology (HPO^4^) as the standardised phenotype vocabulary for recording patient phenotypes, which were entered manually from patient notes by genetic consultants and extracted computationally from patient letters using cTAKES^5^. Patient relatedness was calculated using KING^6^, and any related individuals were filtered out so that this does not skew the probabilities for genetic association studies. This resulted in a subset of 3288 exomes from unrelated individuals.

### Variant call, filtration and phasing

The variant calling and annotation pipeline has been described previously^3^. A total of 973,426 variants were annotated with gnomAD frequencies^7^ and CADD phred score^8^. GnomAD was used as it remains the largest resource for variant population frequency annotation; and CADD owing to its popularity, ability to predict indels and ease of local installation. For gnomAD frequency (GF) annotation, we used both gnomAD allele frequency and estimated homozygote frequency. Estimated homozygote frequency was calculated as:

*homozygote count* × 2 / *allele number*. Variants that did not pass GATK filters, or were not covered by the gnomAD database, were filtered out. Noncoding variants were removed if they are more than 4 bases away from the closest exon. Variants with a missing rate of more than or equal to 20% were discarded (92% mean call rate). Variants in each exemplar gene were phased using Shapeit^9^ for 1000 times. Variant pairs which phased over 75% of the time into the same haplotype in an individual were assumed to be in cis. Accordingly, the analyses of recessive phenotypes would ensure that the filtered variants in a patient are distributed on both alleles of a gene.

### The Phenogenon HPO Goodness of Fit (HGF) Score

Once a gene and a phenotype have been selected for association analysis, variants found on the gene are first binned according to their Gnomad Frequency (GF) and CADD phred score (See Figure 1A and Figure 1B). In the case of dominant Mode of Inheritance (MOI), GF refers to the total allele frequency, while for recessive MOI GF refers to the estimated homozygous frequency. Binned variants are then used to identify patient carriers, and they are separated into two groups for whom the selected phenotype is classified as having the phenotype (Yes) or not having the phenotype (No) (See Figure 1B). The association of the gene and the phenotype within the bin region can then be quantified using Fisher’s exact test (See Figure 1C). Once this is repeated for all binned areas, this can be combined to produce a heatmap (See Figure 1D).

**Figure 1:**
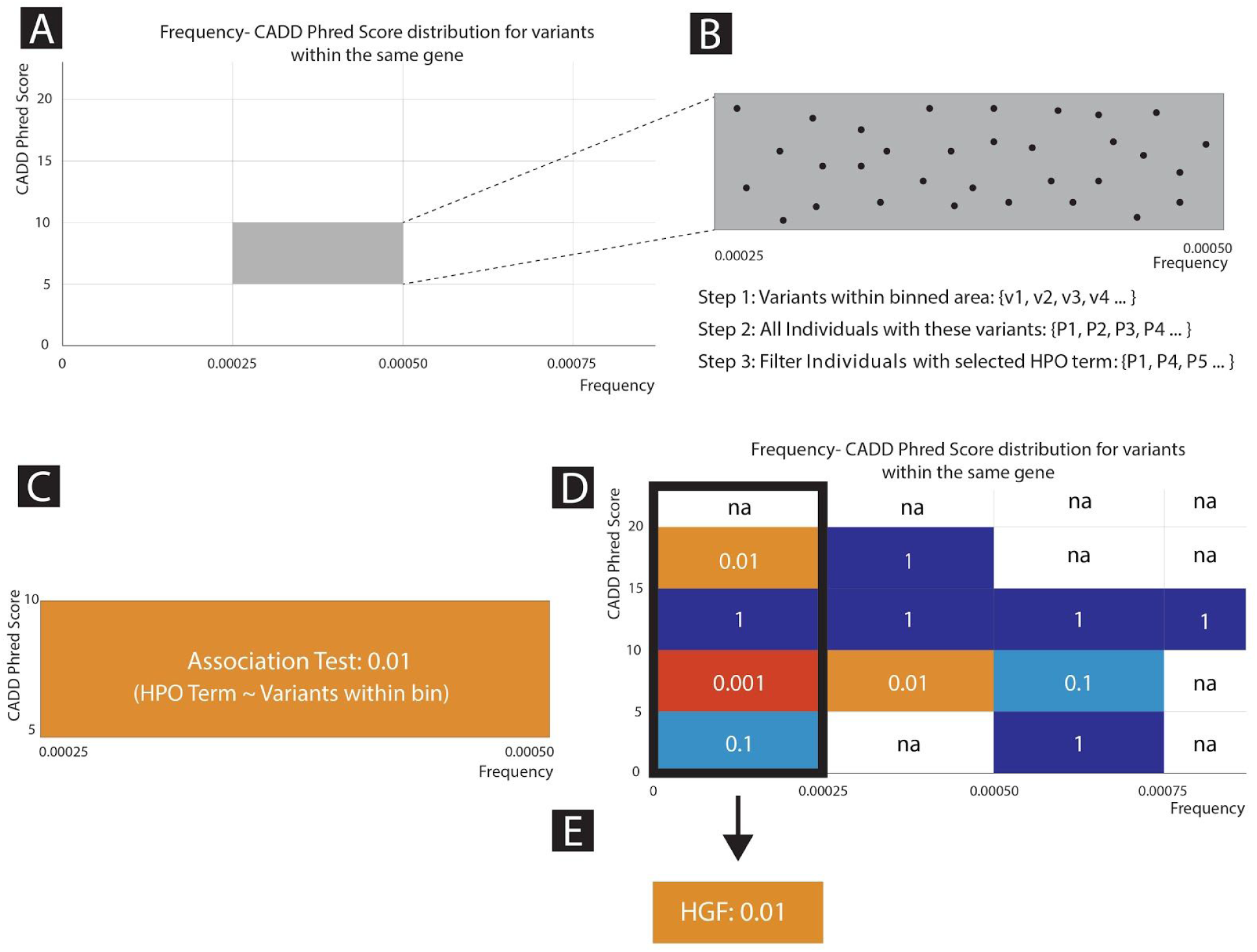
Phenogenon Profiling Workflow. A) The distribution of frequency vs CADD phred score for variants of a single genes are binned according to empirically chosen cut-offs. B) Variants within each binned area are further analysed. Individuals carrying these variants are identified, and then filtered on the basis of whether they have a selected HPO term. C) Fisher’s Exact test is then used to determine the significance of the genotype-phenotype relationship. D) A Phenogenon heatmap is produced using the Fisher Exact P-Values for each binned area. E) Fisher Exact Scores for each of the binned area in the first column are collapsed into a single HPO goodness of fit score (HGF) using a Scaled Stouffer transformation.

Our analyses have shown (described later) that rare variant bins (gnomAD frequency lower than 0.00025) are effective in capturing genotype-phenotype relationships and it is, therefore, favourable to combine the *p* values of the rare variant bins (e.g. combing the first column of the heatmap shown in Figure 1D) with, for example, Fisher’s method. Stouffer’s Z-score method, instead of combining *p* values, combines Z-scores, allowing incorporation of study weights^10^. This gives us an opportunity to assign higher weights to bins with higher CADD phred scores:

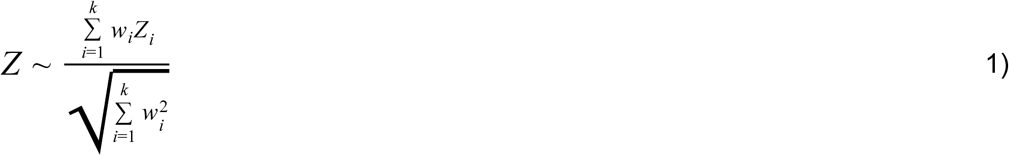

Where *Z_i_* = φ^−1^ (1 − *p_i_*), and φ is the standard normal cumulative distribution function, *p_i_* is the *p* value of the *i*th observation (or bin). We assume that bins with a lower CADD phred score (i.e. lower than 5) are less likely to produce noise and as such can be assigned a low weight. Conversely, bins with a high CADD phred score (i.e. higher than 15) are a more reliable source for evaluating the gene-phenotype association, and therefore deserve higher weights. However, it is still difficult to compare Z scores across genotype – phenotype relationships. For example, if in two relationships there is only one bin that produces the same Z score, where the first relationship has a strong bin with CADD phred window of [0, 5), and the second relationship has a strong bin with CADD phred window of [30, 35), although a low weight (*w_l_*) might be assigned to the bin of the first relationship, and a high weight (*w_h_*) is assigned to the bin of the second, they still yield the same overall Z score, because:

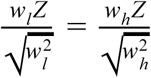
which is not as desired.

To make Z scores comparable across genes, we adjusted formula 1) as follows:

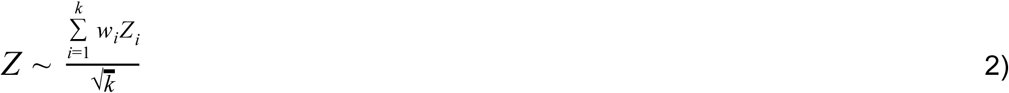

This Z score is converted to a *p* value assuming standard normal distribution. We use the logarithm of the *p* value as the **HPO goodness of fit (HGF) score**.

### Predicting Gene-associated HPO terms

After mapping patients in step 3) in Figure 1, step C - E can be repeated for every HPO term to be tested, and thus produces an HGF score for each HPO term. An HGF cutoff is determined such that HPO terms with an HGF higher than the cutoff are deemed most likely to be associated with the tested genotype. In this study, we used *mean(HGF*) + *stdev(HGF*) as the cutoff, where *stdev* stands for standard deviation.

### Predicting the Mode of Inheritance (MOI)

HPO term prediction can be performed assuming both dominant and recessive MOI. However, in order to identify patients with recessive MOI, a second variant is required for each patient. In this study, the second variant has to meet the lower criteria of a given bin in order to take into account patients carrying compound heterozygous variants. For example, if a patient is selected for a bin because they carries a first variant with GF ∈ [0.00025, 0.0005) and CADD phred ∈ [5, 10), a second variant has to meet the requirement of GF < 0.0005 and CADD phred >= 5. After HPO terms are selected for each MOI, a signal ratio is calculated for each HPO term. This is based on the observation that if a wrong MOI is assumed, the Phenogenon profile tends to produce more bins with low *p* values outside of the first column. The signal ratio can be calculated as:

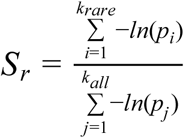

A score for a HPO term can therefore be calculated as:

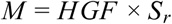

We then used the difference between *M_r_* (when assuming recessive MOI) and *M_d_* (when assuming dominant MOI) for the MOI score:

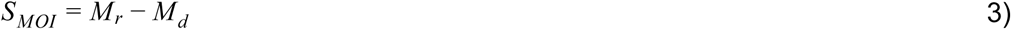

Therefore, a positive *S_MOI_* predicts recessive MOI, and a negative *S_MOI_* predicts dominant MOI. An absolute *S_MOI_* lower than 1.0 can be treated ambiguous.

### The Sequence Kernel Association Test (SKAT)

Single variant testing does not have enough sufficient statistical power to identify an association between a phenotype of interest and a rare variant. Methods were thus subsequently developed to increase power by pooling variants within a given region, typically a gene. Original burden tests treated variants as unidirectional and assumed that pathogenicity increases inversely to variant frequency ^11,12^. The SKAT is a supervised method that performs a joint regression for each variant within a given region^13^. SKAT has more power than burden tests when variants either have variable effect sizes or effects in different directions, i.e. some SNPs in a gene can be protective and some deleterious. The optimal version of SKAT, SKAT-O, was used here as a comparison for the Phenogenon profiling (Supplemental) as it reverts to a burden test if contained variants are unidirectional^14^.

## Results

Based on the number of patients whose genetic causes have been determined, we selected a list of genes for cutoff analysis (Table 1). This is to ensure any positive gene-HPO relationship tested is supported by true knowledge. Their related HPO terms to be tested, the inheritance mode and age of onset were determined primarily based on the resolved patient notes, and published literature. In the following text, we will predominantly use *SCN1A* and *ABCA4* as examples, and results for other genes can be found in the Supplementary materials. Briefly, *SCN1A* encodes Sodium Voltage-Gated Channel Alpha Subunit 1, mutations of which have been linked to epilepsy with divergent clinical severity^15^. The mutations are either dominantly inherited or arise *de novo^16^*. Notably, the majority of *SCN1A* mutations are found in the severe form of epilepsy (severe myoclonic epilepsy in infancy, or SMEI; MIM# 607208), and most are *de novo^15^*.

**Table 1.**
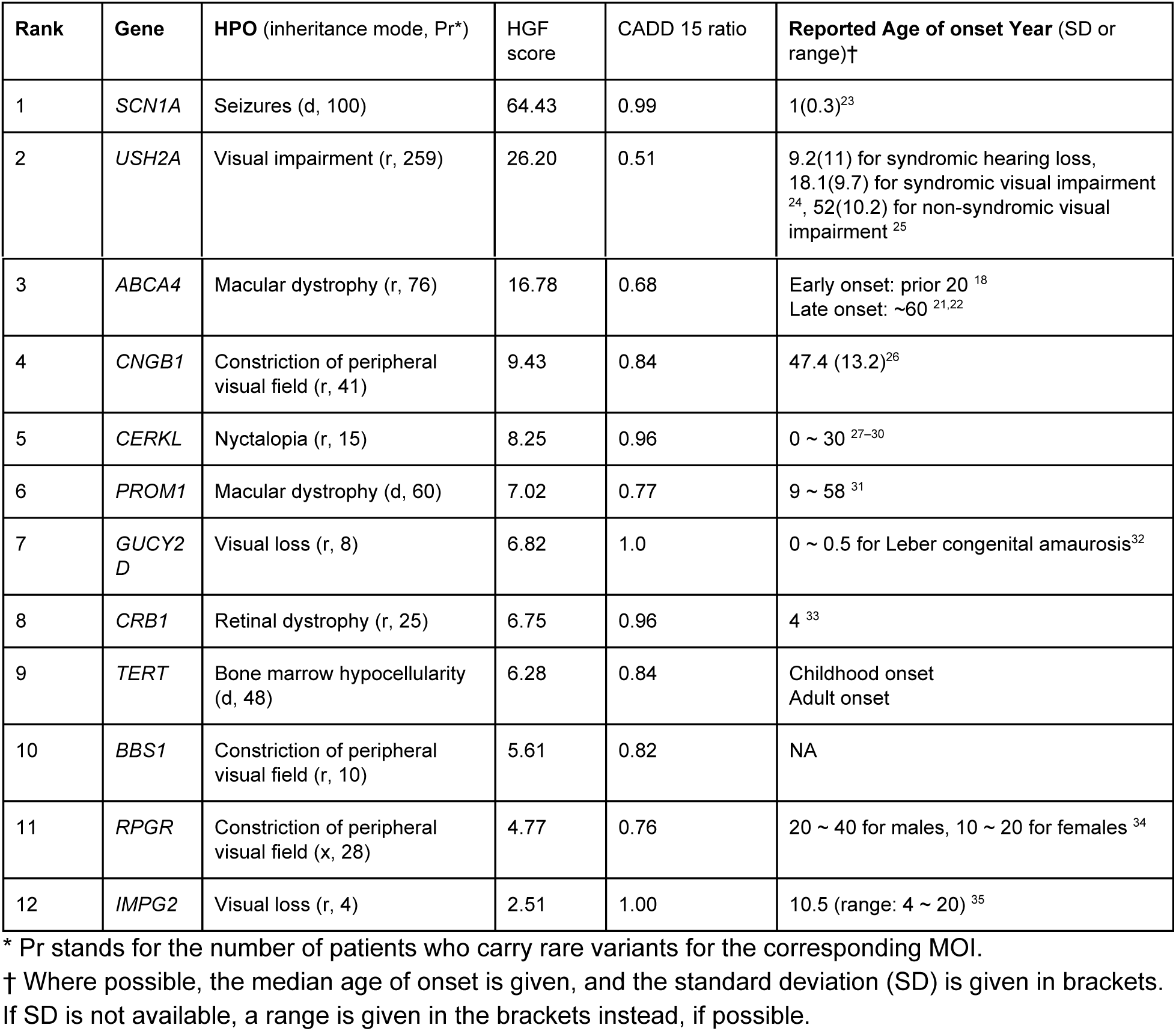
The HPO term with best HGF score per gene in the study and the respective ages of onset where known.

*ABCA4* encodes ATP Binding Cassette Subfamily A Member 4, and its malfunctions are found in patients with retinal abnormalities, including autosomal recessive Stargardt disease, autosomal recessive cone-rod dystrophy, bull’s eye maculopathy, age-related macular degeneration in heterozygous carriers, and autosomal recessive retinitis pigmentosa^17^. Patients with Stargardt disease or retinitis pigmentosa typically have an age of onset prior to 20 years^18–20^, but late-onset Stargardt diseases have also been reported, with age of onset around 60 years^21,22^.

### Predict HPO

We first set out to find how choosing different GF and CADD phred cutoffs may affect variant enrichment for different HPO terms. As shown in Figure 2A, for both *ABCA4* - Macular dystrophy and *SCN1A* - Seizures, bins showing strong association predominantly clustered with variants of GF: 0 ~ 0.00025. From now on, we will denote such variants as rare variants.

**Figure 2.**
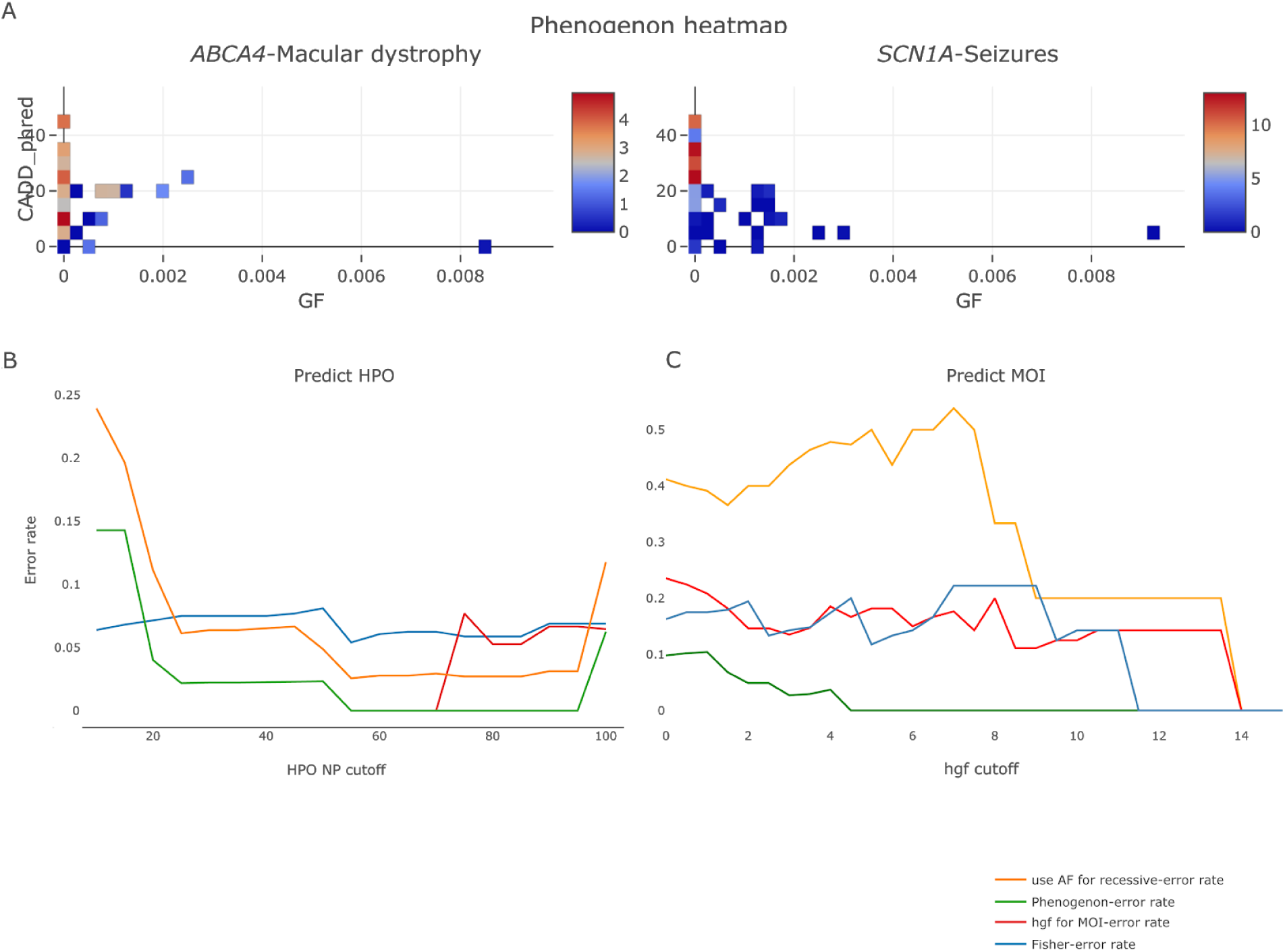
Using Phenogenon to profile gene-HPO relationships, and predict HPOs and mode of inheritance (MOI) for the 12 selected genes. A. Examples of using Phenogenon to profile ABCA4 - Macular dystrophy (HP:0007754) assuming recessive MOI, and SCN1A - Seizures (HP:0001250) assuming dominant MOI. The scales represent – *ln*(*p_i_*) where *p_i_* is the *p* value of the *i*th association test. Note that majority of high value bins are clustered with the rare variant bins. B. Using Phenogenon profiling to predict HPOs. We chose to use HPOs with number of patients higher than ‘HPO NP cutoff’ to assess the performance of the prediction. The staggered lines give the trend of error rates for each prediction model. C. When HPOs are given for each gene, predict MOI. The x axis (HGF cutoff) was used to select HPOs for each gene with HGF score equal to or higher than a given HGF cutoff. The legend is shared for B and C: orange line: use gnomAD allele frequency, instead of estimated homozygote frequency, when assuming recessive MOI; green line: using HGF for HPO association and MOI score for MOI prediction; red line: using HGF for both HPO association and MOI prediction; blue line: using Fisher method to combine *p* values.

We surmise that combining *p* values from the rare variant bins can reflect the strength of genotype-phenotype relationships for rare Mendelian cases. This is indeed the case: Phenogenon correctly predicted HPO terms for which there are at least 55 patients affected (HPO NP >= 55; Figure 2B ‘Predict HPO’, green lines), although as expected, the error rate increased when including rare HPO terms (HPO NP <=20). Interestingly, the error rate increased when HPO NP = 100 (Figure 2B). A closer inspection revealed the source of the mistake as IMPG2, which had only 4 rare variant carriers for the true MOI (recessive). Our model struggled to make correct MOI prediction when less specific HPO terms were included (as shown in Figure 2B, at HPO NP = 100), and correspondingly made wrong HPO prediction when it assumed the wrong MOI.

Notably, combining *p* values using our modified Stouffer’s Z-score method (Phenogenon, green lines in Figure 2B) clearly outperformed Fisher’s method (Fisher, blue lines and bars in Figure 2B), demonstrating the benefit of assigning higher weights to bins with higher CADD phred score.

The HPO term with highest HGF score for each tested gene can be found in Table 1.

### Predicting Mode of Inheritance

A common use case is to ask “which MOI” when a novel gene is under consideration for a patient’s disease. If the gene is more likely to cause recessive MOI whereas the patient’s disease is known to have a dominant family history, then the gene can be assigned with a low priority status.

During the course of this study, we found that when assuming the wrong MOI, the heatmap was more prone to produce low-p-value bins on common variants, a likely source of noise. We used signal ratio (S_r_) to denote how much of such noise we observe from the heatmap: the lower the signal ratio, the higher the noise contributed from common variants. This ratio is incorporated in formula 3 to produce a MOI score: The higher the MOI score (above 0), the more likely a genotype-phenotype relationship is recessive; the lower the MOI score (below 0), the more likely the relationship is dominant. As shown in Figure 2C, using MOI score made fewer mistakes predicting MOI than using HGF alone. The true MOI can be found in Table 1.

### Use estimated homozygote frequency for recessive MOI

Until the release of the gnomAD database, there was no reliable source to estimate variant homozygote frequency, and therefore to date all genotype-phenotype association tools that use population frequency in the model use allele frequency, regardless of MOI in question. We argue that using homozygote frequency when assuming recessive MOI will improve the model performance.

As shown in Figure 2B,C, using gnomAD allele frequency (‘use AF for recessive’, orange lines) when assuming recessive MOI had a poorer performance than using estimated homozygote frequency (‘Phenogenon’, green bars and lines) in predicting HPO and MOI.

### Variance in CADD efficacy

Unlike other tools such as Polyphen and SIFT, CADD does not produce an ordinal value to indicate whether a variant is likely ‘Damaging’, ‘Possibly damaging’ or ‘Benign’. Instead, the authors suggested to use 15 as a cutoff in the task of variant prioritisation. However as shown in Figure 2A: for “*SCN1A* - Seizures”, 15 is sufficient in discriminating potentially pathogenic variants from benign ones; but for “*ABCA4* - Macular dystrophy”, there is no clear cutoff that can separate disease associated and benign variants. This variance may have a random source, or correlate with MOI or age of onset. In order to find the answer, we use CADD phred score 15 as a cutoff to determine variant pathogenicity. As described in Materials and Methods, This produces a CADD 15 ratio with a range of [0, 1]. The closer the ratio to 1, the more efficient to use CADD phred score 15 as a cutoff for variant prioritisation.

When we factored the 12 tested genes according to their MOI, we did not see a significant difference in their CADD 15 ratios (Figure 3, left panel). This might be inconclusive since there was an imbalance of the number of samples in the two groups (dominant MOI: 6; recessive MOI: 37).

**Figure 3.**
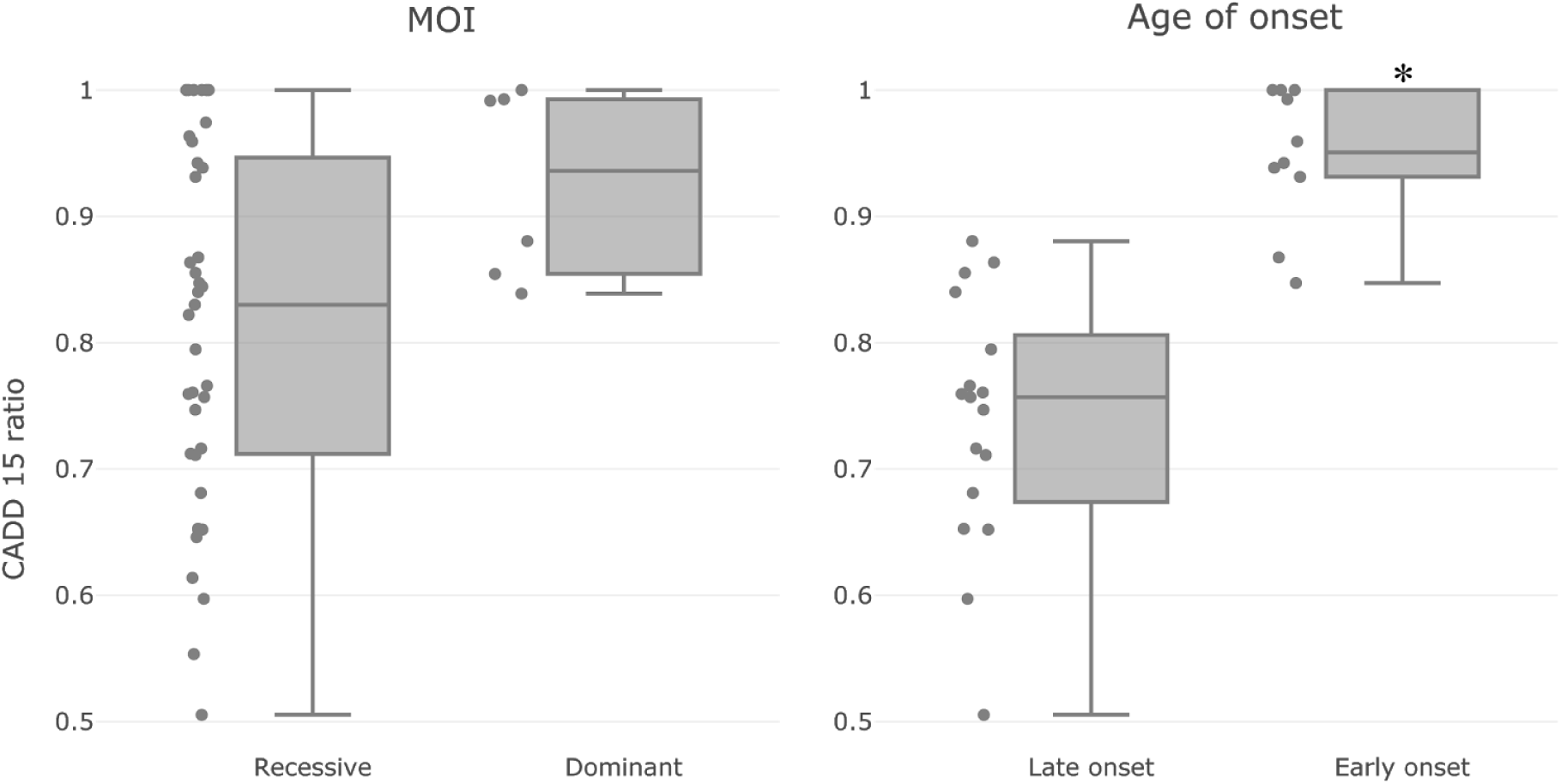
CADD efficacy on genotype-phenotype relationships with different attributes: MOI (left) and Age of onset (right). Each dot represents a genotype-phenotype relation. Briefly, the CADD 15 ratio is calculated to reflect the effectiveness of using *pathogenic* variants only (CADD phred score >= 15) for association test, in comparison with using *non-pathogenic* variants only (CADD phred score < 15). The CADD 15 ratio is normalised to have a range of [0, 1]. The closer the value to 1, the more effective to use CADD phred score 15 as a cutoff for variant prioritisation. There is no significant difference between the dominant and recessive groups of relationships (Mann Whitney U test *p* = 0.04); on the other hand, Early onset relationships tend to have higher CADD 15 ratios than late onset relationships (Mann Whitney U test *p* = 2.6*e*^−5^). Dominant genes: (*SCN1A*, *PROM1*, *TERT*), Recessive genes: (*ABCA4*, *USH2A*, *CNGB1*, *GUCY2D*, *CRB1*, *IMPG2*, *CERKL*, *BBS1*, *RPGR*). Early onset genes: (*SCN1A*, *GUCY2D*, *CRB1*). Late onset genes: (*ABCA4*, *USH2A*, *PROM1*, *CNGB1*, *RPGR*).

Factoring genes according to Age of Onset (AOO) was more challenging, since it was not available in the majority of the sample records. We decided to use literature supported AOO instead as an estimate, and classified those as early onset with AOO likely before the 10th year (*SCN1A, GUCY2D* and CRB1); and those as late onset with AOO likely after the 10th year (*ABCA4, USH2A, PROM1*, *CNGB1* and *RPGR*). Reported AOO can be found in Table 1. As shown in Figure 3, right panel, we found that the early onset genes tend to have a higher CADD 15 ratio than the late onset genes with a Mann Whitney U test *p* value of 2.6*e*^−5^.

### Top ranked genotype-phenotype relationships

Apart from the aforementioned cases, we were also able to uncover other strong genotype-phenotype relationships (Table 2). For instance, *GRHL2* (OMIM: 608576) was known to cause recessive ectodermal dysplasia/short stature syndrome, which involves nail dystrophy^36^, and was correctly linked to Nail dystrophy with recessive MOI by Phenogenon (HGF score: 12.02). *STAT1* (OMIM: 600555) was known to cause dominant or recessive immunodeficiency, and was also correctly linked to Severe combined immunodeficiency, with dominant MOI (HGF score: 11.10). Other examples include *ATG16L1* - Severe combined immunodeficiency (HGF: 9.33) with recessive MOI (known to cause bowel inflammation) and *PMVK -* Nail dystrophy (HGF: 9.03) with recessive MOI (known to cause dominant porokeratosis)

**Table 2.** Top ranked gene - phenotype relations uncovered by Phenogenon

| Gene | Predicted HPO (HGF score) | Known | MOI (predicted / reported)* | Description |
| --- | --- | --- | --- | --- |
| *SCN1A* | Seizures (65.17) | Y | d/d |  |
| *USH2A* | Visual impairment (20.68) | Y | r/r |  |
| *GRHL2* | Nail dystrophy (12.02) | Y | r/r |  |
| *STAT1* | Severe combined immunodeficiency (11.1) | Y | d/dr |  |
| *DNAH9* | Retinal dystrophy (10.13) | N | r/ | *DNAH9* encodes a subunit of an axonemal dynein that mediates the movement of cilia and flagella |
| *ABCA4* | Macular dystrophy (9.65) | Y | r/r |  |
| *ADGRV1* | Rod-cone dystrophy (9.62) | Y | d/r |  |
| *TTN* | Abnormality of the anterior segment of the globe (9.60) | N | d/ |  |
| *ACRBP* | Abnormality of the oral cavity (9.53) | N | d/ | The ACRBP protein is a member of the cancer/testis family of antigens and it is found to be immunogenic, and its expression is found in many tumor types such as bladder, breast, lung, liver, and colon. |
| *VPS13B* | Visual impairment (9.4) | Y | d/r | Known to cause autosomal recessive Cohen syndrome, which also involves retinal dystrophy |
| *SIPA1L3* | Abnormal electroretinogram | N | d/ | Known to cause autosomal recessive congenital cataract; highly expressed in retina |
| *C12orf50* | Neurodevelopmental abnormality (Nail, BM, Oral, Immune) (9.39) | N | r/ |  |
| *ATG16L1* | Severe combined immunodeficiency (9.33) | N | r/ | Known to cause Crohn disease. Produces a protein necessary for autophagy. |

\* MOI: d for dominant, r for recessive, dr if a gene may cause the disease in either dominant or recessive MOI.

We also found links between genotype and phenotype not reported before. *SIPA1L3* (OMIM: 616655), a gene associated with recessive congenital cataract, was found to have a high HGF score on Abnormal electroretinogram (HGF: 10.66) and Rod-cone dystrophy (HGF: 7.45), with dominant MOI. Although no *SIPA1L3* mutations have been reported with relevance to retinal dystrophy, it was found to be highly expressed in *Xenopus laevis* retina^37^, indicating a link between this gene and retina disorders. *ATG16L1*, known to cause Crohn disease, was linked to ‘Severe combined immunodeficiency’. In fact, Crohn disease was recently suggested as a result of immunodeficiency^38^, and since *ATG16L1* encodes a protein necessary for autophagy, this gene could as well be a genetic cause of the patients that have ‘Severe combined immunodeficiency’ in UCLex.

## Discussion

Aggregated databases of high throughput sequencing data of large numbers of patients, from varying ethnicities with many phenotypes, are increasingly the norm. This is becoming mandatory as large sample sizes are needed to have sufficient statistical power for finding causative variants of rare diseases. The UCLex database consists of genotype data and phenotype data with mixed levels of qualities. For example, of 3288 individuals in our analysis, 2423 had only one recorded HPO term, the majority of whom had Dementia (1039). Given that dementia is a disease of old age, patients would be expected to present with comorbidities; but this is not reflected in the database. Excluding the one-HPO individuals, the median number of HPO terms for an individual is 4, with a standard deviation of 3.6.

Another challenge of large and mixed cohort is the ambiguity between absence of evidence and evidence of absence. Since for rare phenotypes most of the time absence of evidence is equivalent to evidence of absence, collapsing both cases to evidence of absence works well. However, for less rare phenotypes such as Cataract (HP:0000518), it is often miss-reported in patients whose primary disease is not at the eyes, and hence its presence is falsely correlated with other eye diseases such as retinal dystrophy. To make it worse, diseases with much earlier onset than retinal dystrophies will be less likely to allow the patient to develop cataract at the time of diagnosis. We, therefore, removed cataract-related terms from our analysis.

Despite the very different levels of phenotype annotations and inconsistent genomic sequence captures in the UCLex samples, using our new tool, Phenogenon, we were able to uncover genuine relationships between genes and phenotypes. Specifically, it was achieved with reference to variant population frequencies (gnomAD allele frequency or estimated gnomAD homozygote frequency) and a variant scoring tool (CADD phred).

We tested a series of variant population frequency bin sizes from 0.001 to 0.0001, and Phenogenon performance optimised at 0.00025, without any substantial improvement by further reducing the bin size (data not shown).

Nine out of twelve testing genes may cause recessive MOI; similar odd ratio is also observed in RetNet (90 out of 118). In fact, most of the rare Mendelian cases, especially those with an early age manifestation, are recessive. Modelling recessive MOI proved difficult since 1) phasing variants on such a large dataset is challenging and 2) the requirement on prediction accuracy of both alleles pathogenicity is more demanding than the prediction of one allele. Additionally, in order to maintain a reasonable false positive rate of our model, we removed the majority of the non-coding variants where CADD has a much lower power in pathogenicity prediction. If the molecular cause of a disease frequently involves combinations of an intronic variant and a coding variant, Phenogenon will mistakenly produce a dominant MOI score. Therefore an efficient phasing tool for large data set of exome/genome sequencing data, and a pathogenicity tool with an improved power of pathogenicity prediction on non-coding variants will greatly benefit recessive MOI modelling.

Phenogenon can also be used in gene prioritisation when a combination of phenotype terms are under consideration. When we took the combination of ‘Rod-cone dystrophy’ (HP:0000510) and ‘Hearing impairment’ (HP:0000365) as an input, the top two genes predicted by Phenogenon were *USH2A* (HGF: 9.53) and *ADGRV1* (HGF: 8.31), both are known to cause Usher syndrome that affects both visual and hearing systems. However, an issue accompanying such approach is a much-reduced size of affected patients, which will reduce Phenogenon prediction power. The utilisation of the HPO nomenclature by the 100,000 genomes project means that this combinatorial analysis will be of increasing value as genetics is phased into routine NHS practice.

## Acknowledgements

We thank Lucy Withington, Stephanie Halford and Suzanne Broadgate for their indispensable help during the development of the work described in this paper. This work is funded by UKRP Fighting Blindness.

## Supplementary

**Supplementary Figure 1.**
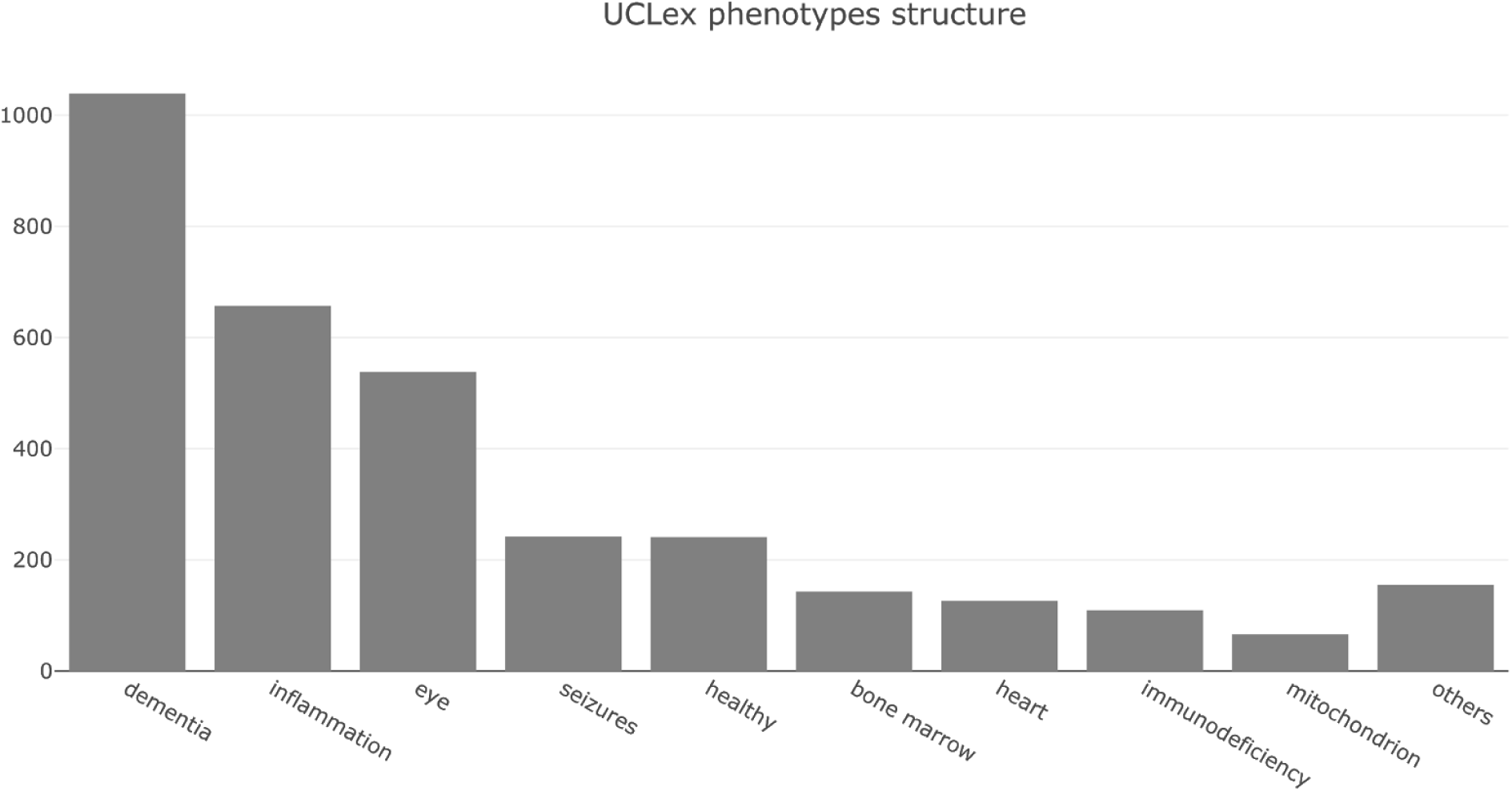
UCLex phenotypes structure. We grouped all the unrelated patients in this study into major phenotype categories according to their registered HPO terms. Note that the largest two groups (dementia and inflammation) are not likely to have many Mendelian cases. The third and fourth most popular groups (eye and seizures) have the most known Mendelian cases such as *ABCA4* - Macular dystrophy and *SCN1A* - Seizures.

### Comparison to the Optimal Sequence Kernel Association Test

In order to test the robustness of our analysis, we applied a regression approach taken in the Optimal-Sequence Kernel Association Test (SKATO)^13^ on the 12 exemplar genes (Supplementary Figure 2). The strong association between *SCN1A* and epilepsy was replicated, as was *ABCA4* and macular dystrophy. However, SKAT did not match the performance of Phenogenon when testing retinal dystrophy (Table 3). Additionally SKAT was not able to stratify genes by age of onset (Data not shown).

**Table 3.** SKATO test statistics on a selection of the 12 exemplar genes

| Gene | Phenotype | SKAT-O | Case maf (n) | Ctrl maf (n) |
| --- | --- | --- | --- | --- |
| *SCN1A* | Epilepsy | 1.1423e-20 | 0.00066 (242) | 1.2e-04 (2991) |
| *ABCA4* | Retinal Dystrophy | 0.00744 | 0.00029 (434) | 0.00016 (2820) |
| *USH2A* | Retinal Dystrophy | 0.02241 | 0.00018 (434) | 0.00014 (2820) |
| *ABCA4* | Macular Dystrophy | 2.5888e-12 | 0.00058 (70) | 0.00017 (3179) |

**Supplementary Figure 2.**
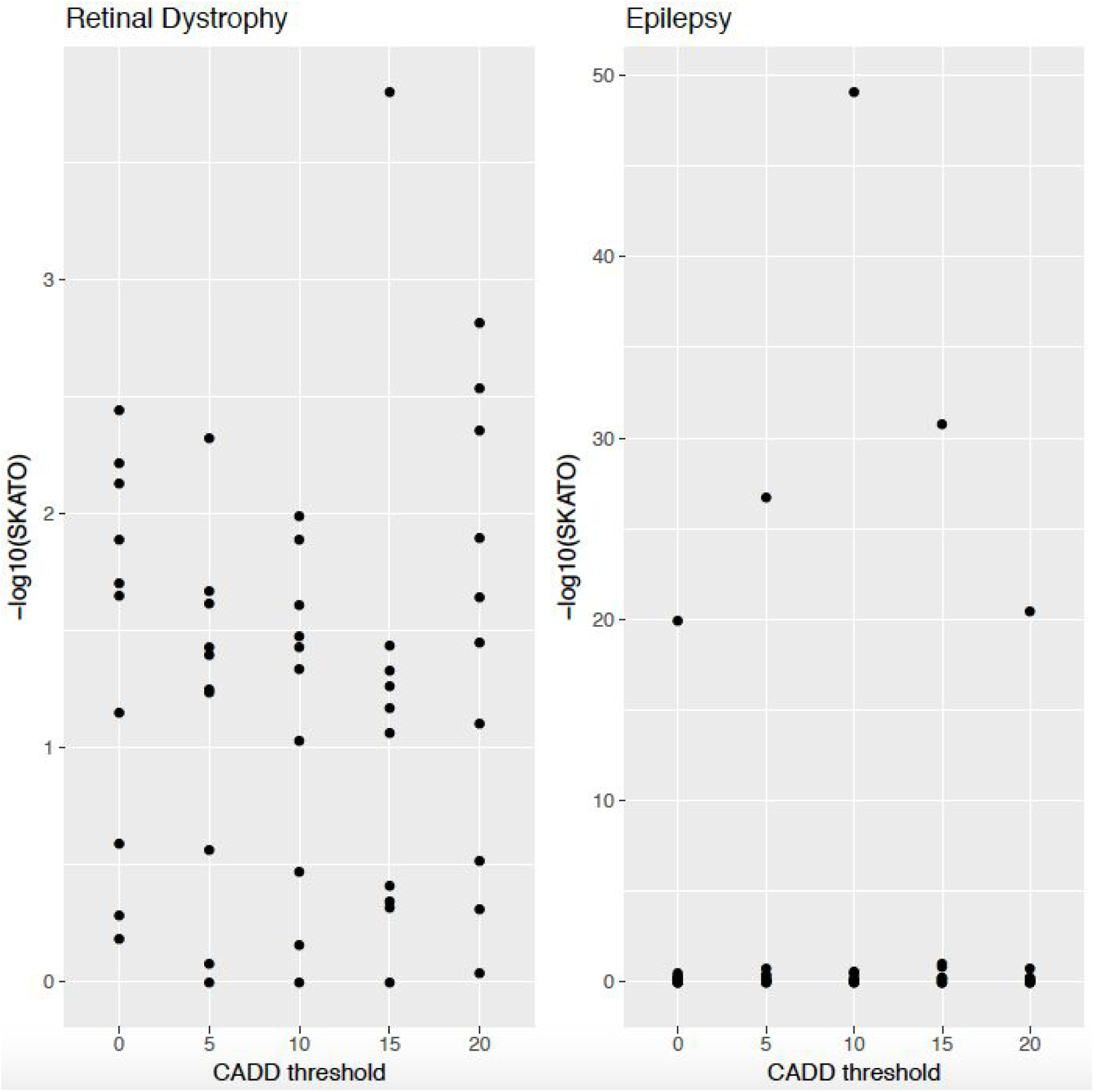
Assessing the impact of varying minimum CADD phred score on SKAT-O p-value for the Retinal Dystrophy and Epilepsy cohorts. Each dot represents a p-value for each of the 12 principal genes that contains variants post filtering.

